# Detection of High-Frequency Oscillations from Intracranial EEG Data with Switching State Space Model

**DOI:** 10.1101/2024.05.01.592107

**Authors:** Zeyu Gu, Shihao Yang, Zhongyuan Yu, Feng Liu

## Abstract

High Frequency Oscillations (HFOs) is an important biomarker that can potentially pinpoint the epileptogenic zones (EZs). However, the duration of HFO is short with around 4 cycles, which might be hard to recognize when embedded within signals of lower frequency oscillatory background. In addition, annotating HFOs manually can be time-consuming given long-time recordings and up to hundreds of intracranial electrodes. We propose to leverage a Switching State Space Model (SSSM) to identify the HFOs events automatically and instantaneously without relying on extracting features from sliding windows. The effectiveness of the SSSM for HFOs detection is fully validated in the intracranial EEG recording from human subjects undergoing the presurgical evaluations and showed improved accuracy when capturing the HFOs occurrence and their duration.

## I. INTRODUCTION

High frequency oscillations (HFOs) can be detected from the brain electrophysiological activities through intracranial EEG (iEEG) recordings captured by electrodes with multiple contacts. These oscillations are generally classified into two types based on frequency range: Ripples (80–250 Hz) and fast ripples (250–500 Hz) [1][2][3]. HFOs are transient, low-amplitude fluctuations of varying frequencies, shapes, and pathophysiological origins, and recent studies have identified them as biomarkers for identifying epileptogenic zones. For drug-refractory epilepsy patients, surgical resection of the identified seizure irritative zones can offer favorable surgical outcome [4][5].

However, due to the variability of the morphology of HFOs, precise detection is an important yet challenging task. The oscillations are short-lived, nonstationary events with low amplitude, indicating a variety of neural activities [1]. Moreover, the iEEG signal can be contaminated by noise from environmental and physiological sources. Additionally, bandpass filters can erroneously detect transient events like sharp waves and interictal epileptiform discharges. Finally, the morphological and spectral diversity of HFOs, plus their variability both within and between subjects, complicates their detection accuracy [6].

Machine learning techniques have effectively been applied to the categorization of HFOs, beginning with methods that rely on manual feature engineering for each HFO event. These methodologies included linear discriminant analysis, support vector machines, decision trees, and clustering [7]. However, these prior deep learning methods for HFO detection have been predominantly supervised, and require human-annotated labels, thus constraining their broader application due to the necessity of expert labeling. In addition, it is time-consuming and prone to human errors.

Neural time series data often exhibit non-stationary behavior with abrupt dynamic shifts and transient activities. State space models (SSMs) are effective for modeling clear trends and cycles in time series. Given the presence of regimes (underlying different brain states) of the EEG time series data, switching state space models (SSSMs) are widely to identify the underlying switching patterns [8][9][10]. The SSSMs offer a more direct approach to characterize time series data with switched regimes by employing a set of sub-SSMs, each representing different dynamics, and toggling between them based on a probabilistic mechanism. However, finding precise state inference and parameter estimation for these switching models is computationally challenging and often not feasible with exact methods.

Recently, a new SSSM inference paradigm was introduced by He et al. [8], which leverages state inference solutions within the generalized expectation-maximization algorithm to estimate model parameters for the switching process, which it is particularly suitable for neural signal analysis. Inspired by He et al. [8], we propose to leverage SSSM for identifying HFOs. The advantage of using SSSM is its capability of identifying HFOs instantaneously without relying on feature extraction within an epoched iEEG window. The main contributions of our work in this task are as follows:

- We applied the novel method proposed by He et al. [8] to the problem of detecting HFOs in the iEEG data.
- We evaluate our model using the Zurich iEEG HFO Dataset [11], demonstrating its ability to achieve more accurate results compared to the original event annotations.
- Our approach is unsupervised, eliminating the need for manual labeling and significantly reducing both workload and potential labeling errors.

## II. METHOD

In this section, the switching state-space oscillator models and Variational Bayesian (VB) learning proposed by He et al.[8] are detailed. An SSSM typically comprises two sets of states: the continuous-valued states and the discrete-valued states. Within a model containing *M* Gaussian SSMs, these continuous-valued hidden states are labeled as 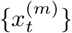 with *m* ∈ {1, …, *M*}, the discrete-valued hidden states of the SSSM are denoted as {*s*_*t*_} where *s*_*t*_ ∈ {1, …, *M*}, and the real-valued observations are denoted as {*y*_*t*_}. In the context of analyzing HFOs, we use two underlying SSMs characterizing non-HFO and HFO time sequences in the SSSM.

### A. Switching state-space models

The SSSM is a sophisticated statistical framework utilized for modeling non-stationary time series data. It features a set of latent variables that transition among distinct states, with each state possessing its dynamic behavioral characterization. The model dynamics are encapsulated by the state transition equation:

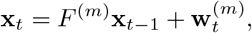

where **x**_*t*_ denotes the state at time *t, F* ^(*m*)^ is the state transition matrix, and 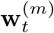 signifies the process noise. The observed data are related to the state variables via the measurement equation:

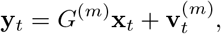

with **y**_*t*_ representing the observed data, *G*^(*m*)^ the observation matrix, and 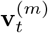 the observation noise. The superscript *m* represents the *m*-th state equation regime. Within the Bayesian inference paradigm, the model employs a switching mechanism governed by a discrete latent variable *s*_*t*_, which determines the appropriate regime at each juncture. This switching mechanism enables the SSSM to adaptively mirror abrupt shifts in the data-generating process, thereby providing a potent analytical instrument for deciphering complex temporal patterns amidst stochastic volatility.

### B. Switching state-space oscillator models

Oscillatory patterns are pervasive in neural signals, necessitating their modeling with a high degree of accuracy and precision to detect any brief oscillatory events. In this context, we utilize a novel state-space model specifically tailored for the task of capturing oscillations [8][12]:

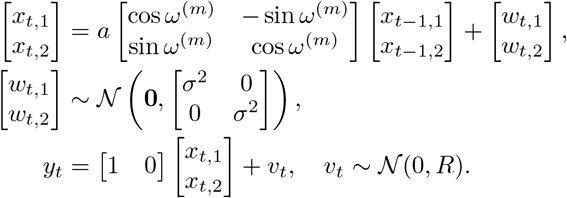

The model can be interpreted as a phasor that rotates around the origin in the complex plane with a frequency *ω*^(*m*)^ (in radians). The constant *a* represents the damping factor, which dictates the rate at which the oscillations diminish over time. We will refer to this model as an oscillator model. This model is used to identify the HFO and non-HFO time sequences.

### C. Variational Bayesian learning

In this part, we will detail the variational Bayesian learning as an instance of a generalized EM algorithm. Previous work has proposed a variational approximation of the posterior of hidden states. One possible solution is to employ the variational Bayes technique to approximate the true posterior of the hidden states [8][13].

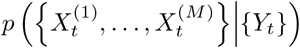

The idea is to operate within a subspace of tractable posterior probability distributions defined over the hidden states, and to choose the optimal approximation,

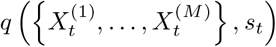

based on a lower bound on the marginal log-likelihood of observations.

Two candidate models are constructed, parameterized by Θ = {*α*^(*m*)^, *ω*^(*m*)^, (*σ*^2^)^(*m*)^} for *m* ∈ {*δ, ϵ*}. Preprocessing of the raw data through a filtering phase to ensure clean input for the algorithm. An iterative process involving:

1. Initialize hidden states 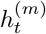 and variational summaries 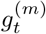 under current parameters.

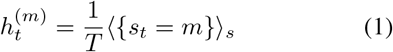

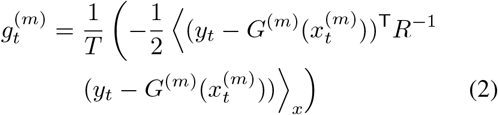

The temperature *T* anneals during the optimization process, affecting the exploration of the solution space and convergence to the optimal approximate distribution [14]. It is updated by the decay function *T*_*i*+1_ = (*T*_*i*_ + 1)*/*2, facilitating a gradual decrease in entropy of the variational free energy.
2. In the algorithm, we first compute the hidden states 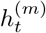 using the forward-backward algorithm on a Hidden Markov Model (HMM), where the observation probability is given by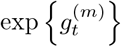 . Subsequently, we calculate the variational summaries 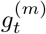 by applying Kalman filtering/smoothing on an SSM, with the observation noise covariance determined by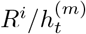 . This sequence of operations refines the state estimates.
3. Parameter updates upon stabilization to Θ^*i*+1^.
4. Convergence check using the negative variational free energy ℱ(*q*, Θ):

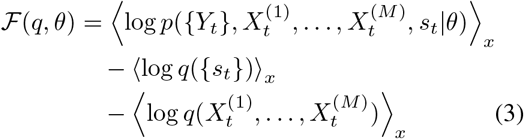

where

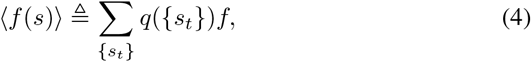

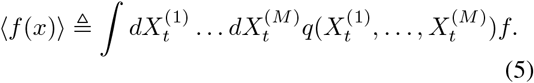

Ultimately, we acquire the model’s output parameter set 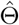, which comprises the means 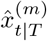, covariances 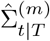,and state probabilities 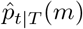 across *M* states. Based on these outputs, we generate a binary variable image to represent the segmentation of HFOs, which is then compared to the annotation of events [13].

## III. RESULT

### A. Datasets

We evaluated our model using the Zurich iEEG HFO Dataset [11]. This dataset contains samples of long-term invasive recordings (iEEG) from 20 patients who subsequently underwent epilepsy surgery. The unique nature of this dataset, with its specific focus on epilepsy patients, allows us to demonstrate the generalizability and applicability of our model in a specialized clinical context.

In the Zurich iEEG HFO Dataset, there are 20 subjects aged between 17-52 years (14 females and 6 males), with detailed intracranial EEG data. Intracranial recordings were acquired at a 2000 Hz sampling frequency with an ATLAS recording system (0.5-1000 Hz pass-band, Neuralynx, www.neuralynx.com) and down-sampled to 2000 Hz for HFO analysis. Sleep scoring was performed based on scalp EEG, electrooculogram, EMG, and video recordings. We chose time intervals at least three hours apart from epileptic seizures to eliminate the influence of seizure activity on our analysis. For our analysis, we exclusively focused on data from two patients to illustrate the effectiveness of the proposed method.

### B. Experimental Setting

For EEG spectrogram analysis, we employed the multitaper method with specific parameters: a sampling frequency (FS) of 2000 Hz, a frequency range limited between 20 and 250 Hz, and a time-half bandwidth of 2. The number of tapers was set to be 3. We chose window parameters with a size of 0.04 seconds and a step size of 0.001 seconds. This method allowed for a detailed and nuanced analysis of the EEG data, ensuring high fidelity in frequency resolution and temporal precision.

We modeled non-HFOs (*δ*) and HFOs (*ϵ*) observed in iEEG recordings using Gaussian SSMs of the following form. The Gaussian SSM parameters, which include non-HFOs 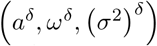 and HFOs 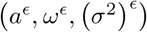, along-side the observation noise variance *R*, were initialized to learn the optimal model parameters using the generalized Expectation-Maximization (EM) algorithm. The initialization was achieved by fitting two oscillators to the EEG time series, assuming the presence of both types of oscillations throughout the duration. This fitting utilized a standard EM algorithm with parameters set based on our prior knowledge of the typical frequencies of these oscillations [8]. The parameters were set as follows: for non-HFOs, 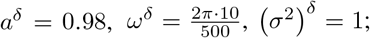 for HFOs, 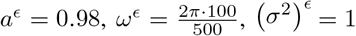 and the observation noise variance, *R=*1

Initial states were assumed as zero-mean white noise with a variance of 3 and were not updated during the iterations. The EM algorithm was run for 50 iterations to establish initial parameter estimates for the Gaussian SSMs within the switching state-space models. The parameters obtained after these iterations were used in the switching inference algorithms, to infer segmentation [15].

### C. Results

Fig. 2. presents a detailed comparison of segmentation performance, specifically illustrating from top to bottom the spectrograms, raw data traces, SSSM algorithm segmentation results, and manual event annotations. The segmentation outcomes from the SSSM and the manual annotations, are demarcated along the y-axis, where 0 and 1 indicate non-HFO and HFO events, respectively.

**Fig. 1.**
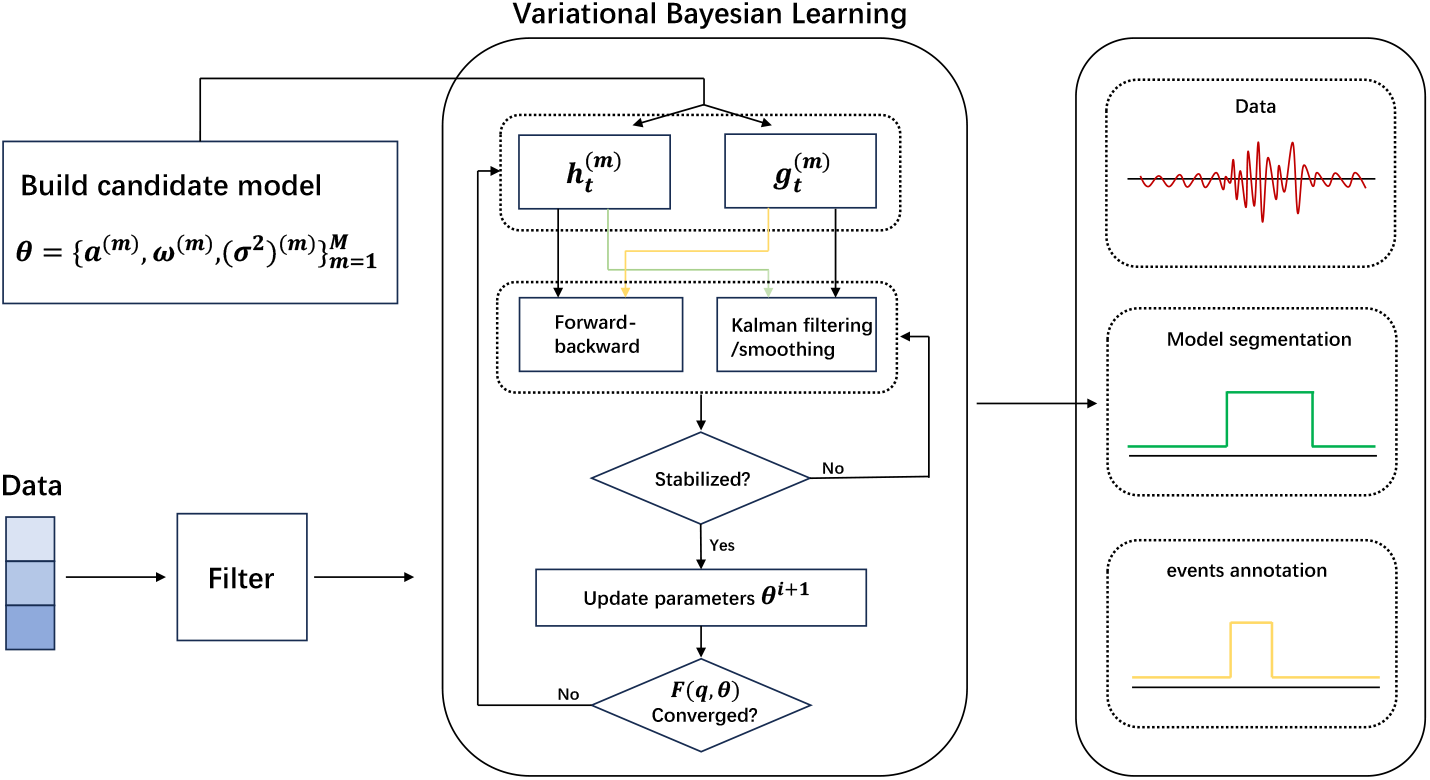
A schematic representation of the Variational Bayesian Learning model is depicted. The procedure initiates with the input of raw data, which subsequently undergoes a filtering stage. Post filtration, the data is introduced into the Variational Bayesian Learning framework, facilitating the construction of a candidate model. The model parameters are recursively estimated, employing Kalman filtering/smoothing alongside a forward-backward algorithm. This iterative process involves stability checks and parameter updates, transitioning from *θ*^(*i*)^ to *θ*^(*i*+1)^. Iteration persists until the convergence criterion *F* (*q, θ*) is satisfied.

**Fig. 2.**
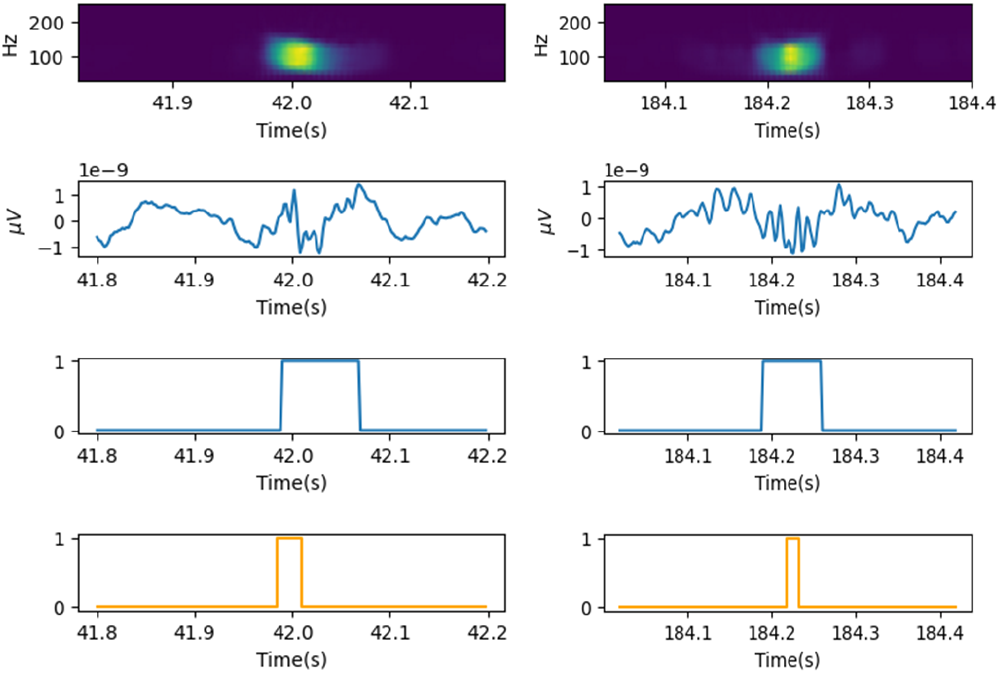
The figure illustrates two HFO detection examples. From top to bottom, the subplots include a spectral map, raw data trace, SSSM algorithm segmentation, and manual event annotations.

The SSSM method successfully segments HFO events, with the segmented areas encompassing the high-frequency components visible in the spectrograms. Notably, our segmentation differs from the annotated events; it does not rely on pre-labeled data for learning but rather models the data directly for analysis. This approach ensures that the segmentation is not biased by pre-existing labels but is instead a true representation of the underlying data. Observations from the actual data indicate that our segmented regions are more accurately aligned with the true events compared to the annotations. We tested the SSSM on two patients of Zurich iEEG HFO Dataset [11], and the performance comparison with Support Vector Machine (SVM), Random Forest (RF), and Deep Neural Network (DNN) is given in Table 1.

**TABLE I.** Model Performance Evaluation

|  | Patient 1 |  |  |  | Patient 2 |  |  |  |
| --- | --- | --- | --- | --- | --- | --- | --- | --- |
|  | Precision | Accuracy | Recall | F1-score | Precision | Accuracy | Recall | F1-score |
| SVM | 0.674 | 0.658 | 0.702 | 0.687 | 0.642 | 0.623 | 0.659 | 0.648 |
| RF | 0.331 | 0.569 | 0.605 | 0.458 | 0.598 | 0.574 | 0.586 | 0.583 |
| DNN | 0.714 | 0.781 | 0.753 | 0.732 | 0.705 | 0.769 | 0.762 | 0.738 |
| SSSM | 0.863 | 0.814 | 0.827 | 0.816 | 0.835 | 0.823 | 0.856 | 0.824 |

This method demonstrates its capability to segment both non-HFO and HFO events in an unsupervised way. It offers an instantaneous and precise interpretation of the data without relying on extracting features from the epoched windows.

## IV. CONCLUSIONS

In this paper, we validated the switching state space model for the detection of HFOs in the iEEG data on patients with epilepsy. This method adeptly models transient neural high frequency burst activities. As demonstrated on the data from epilepsy patients, the proposed SSSM can accurately segment HFOs. The SSSM notably surpasses the benchmark algorithms in detection accuracy. Given that the state space model is inherently univariate, future research can be designed leveraging multichannel iEEG data, exploring the spatial-temporal data structures for the HFOs detection.

